# Pandora: A Tool to Estimate Dimensionality Reduction Stability of Genotype Data

**DOI:** 10.1101/2024.03.14.584962

**Authors:** Julia Haag, Alexander I. Jordan, Alexandros Stamatakis

## Abstract

**Motivation:** Genotype datasets typically contain a large number of single nucleotide polymorphisms for a comparatively small number of individuals. To identify similarities between individuals and to infer an individual’s origin or membership to a population, dimensionality reduction techniques are routinely deployed. However, inherent (technical) difficulties such as missing or noisy data need to be accounted for when analyzing a lower dimensional representation of genotype data, and the intrinsic uncertainty of such analyses should be reported in all studies. However, to date, there exists no stability assessment technique for genotype data that can estimate this uncertainty.

**Results:** Here, we present Pandora, a stability estimation framework for genotype data based on bootstrapping. Pandora computes an overall score to quantify the stability of the entire embedding, infers per-individual support values, and also deploys a *k*-means clustering approach to assess the uncertainty of assignments to potential cultural groups. Using published empirical and simulated datasets, we demonstrate the usage and utility of Pandora for studies that rely on dimensionality reduction techniques.

**Availability and Implementation:** Pandora is available on GitHub https://github.com/tschuelia/Pandora.

**Supplementary information:** Supplementary data are available online.

## 1 Introduction

Dimensionality reduction techniques such as Principal Component Analysis (PCA) and Multidimensional Scaling (MDS) are routinely deployed in numerous scientific fields as they facilitate data visualization and interpretation. Both methods project high-dimensional data to lower dimensions, while preserving as much variation, and thus information, as possible. Ever since the first application of PCA to genetic data almost half a century ago [14], PCA as well as alternative dimensionality reduction techniques have been widely used in modern population genomic analyses, including ancient DNA studies, to draw conclusions about population structure, genetic variation, or demographic history. For example, Hughey et al. [10] found that the Minoan civilization did not originate from Africa, relying on PCA results. However, due to common challenges such as missing data and noise, the deployment of dimensionality reduction techniques to analyze ancient DNA or conduct contemporary population genetics studies is controversial. For instance, Yi and Latch [26] demonstrated that missing data can bias PCA analyses. Thus, quantifying the intrinsic uncertainty is pivotal to proper and cautious result interpretation. For example, the relative location of a population or individual in a PCA/MDS embedding determines its relation to other populations or individuals. If there exists a substantial uncertainty with respect to the projection location for some, or even all individuals in a study, the respective conclusions may potentially be erroneous and hence misleading. To the best of our knowledge, there currently exists no method to quantify the inherent uncertainty of dimensionality reduction techniques in either ancient or contemporary population genetic studies. To address this challenge, we introduce Pandora, an open-source tool that estimates the uncertainty of PCA and MDS analyses of population genetics and ancient DNA genotype datasets. Pandora estimates the uncertainty via bootstrapping over the SNPs. In addition to an overall, global stability score, Pandora also estimates the stability of a subsequent *k*-means clustering based upon the computed embeddings. Both stability scores have values that range between zero and one. Higher values indicate a higher stability and thus a lower uncertainty with respect to the computed embedding for the respective dataset. Beyond this, Pandora also infers dedicated per-individual bootstrap support values for all individuals in the dataset. These support values indicate whether an individual’s location in the lower-dimensional embedding space is stable across bootstrapped embeddings, or, if the individual should be considered as being “rogue” when its projections between distinct bootstrapped embeddings differ substantially. For such a “rogue” individual, any conclusion concerning, for instance, the assignment to a specific (sub-)population should be carefully (re-)considered.

Pandora is implemented in Python3 and is available open-source on GitHub https://github.com/tschuelia/Pandora. It can be used via the command line or as a Python library. A thorough documentation of the tool, including usage examples, is available at https://pandorageno.readthedocs.io.

## 2 Methods

Figure 1 depicts our algorithm to estimate the stability of a genotype dataset under dimensionality reduction via bootstrapping. We initially generate *N* bootstrap replicates by randomly sampling from the set of Single Nucleotide Polymorphisms (SNPs) in the input genotype data *with* replacement. Then, for each bootstrap replicate, we perform a dimensionality reduction using either PCA or MDS, depending on the user’s choice. Based on the computed embedding replicates, we subsequently compute the aforementioned three stability values: the overall Pandora stability (PS), the per-individual Pandora Support Values (PSVs), and, *after* applying *k*-means clustering, the Pandora Cluster Stability (PCS).

**Figure 1:**
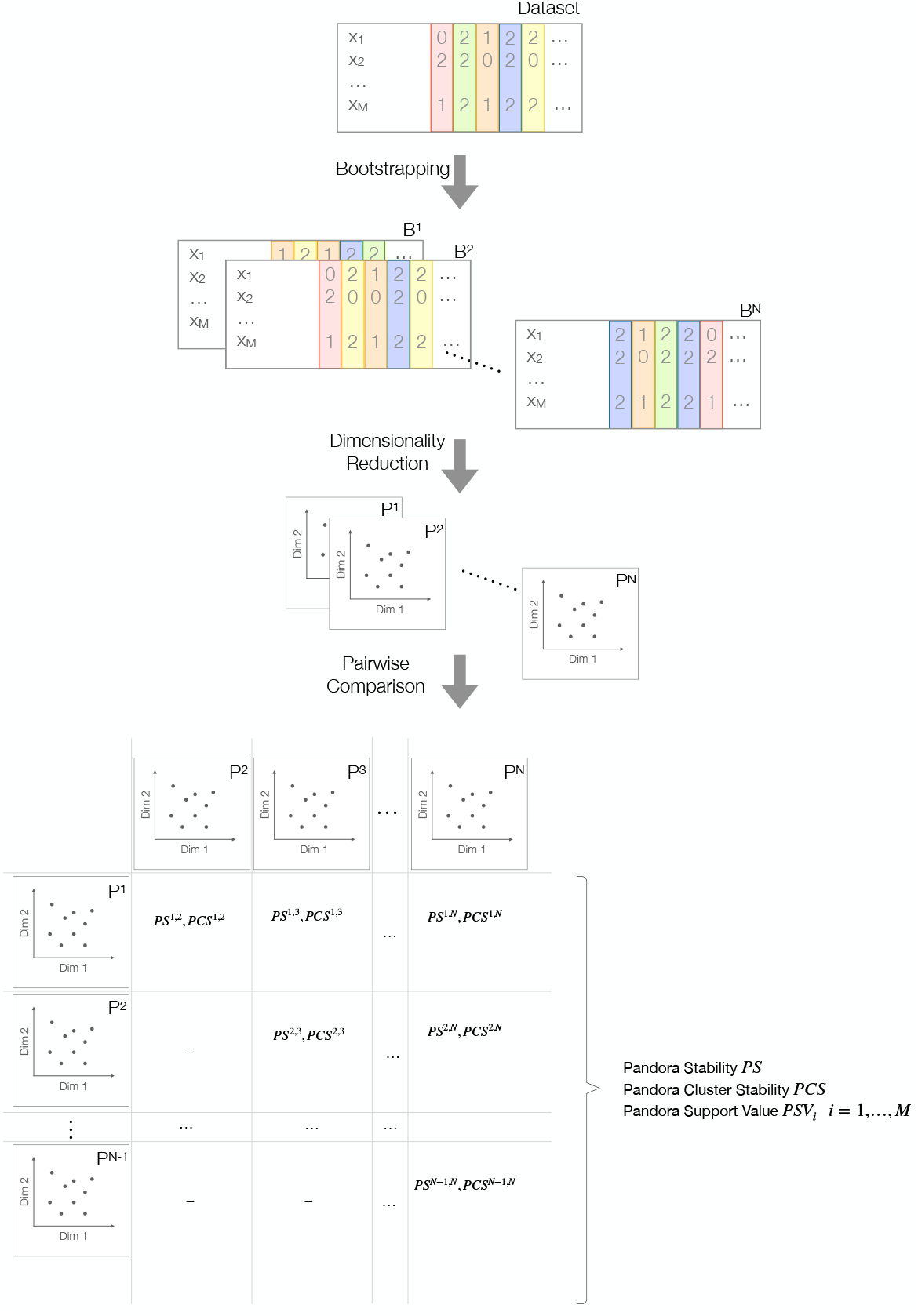
Schematic overview of the bootstrap-based stability analyses for genotype data, as implemented in our Pandora tool.

In the following, we briefly outline the fundamentals of PCA and MDS, and provide information on the respective implementations we use in Pandora. We also describe Pandora’s bootstrap procedure in more detail, focusing on the pairwise comparison of two embeddings, as well as the computation of the PS, PCS, and PSVs values.

Henceforth, the term ‘population’ refers to its biological rather than to its statistical interpretation. We refer to a genotype data sequence in the input data as a single individual and, in general, each individual is associated to a population (e.g., “Greek”).

### 2.1 Genotype Data

Pandora was specifically designed for stability analyses of dimensionality reduction on genotype data. This genotype data consists of sequences of SNPs, that is, the genetic variation between individuals. We further denote the number of sequences, that is, the number of individuals, in the dataset as *M* and the number of SNPs as *K*. Note that Pandora, following the EIGENSOFT [17] file specification, assumes that data are biallelic. In this case, a specific locus in the maternal and paternal genomic data can be either one of two alleles *a* or *A*. Consequently, the genotype of an individual can take one of four states: *aa, aA, Aa*, or *AA*. This genetic information is encoded for each SNP and each individual of a genotype dataset under study. Note that the states *aA* and *Aa* are typically encoded using the same character, since the resulting genotype is independent of the allele order. Finally, a dedicated special character denotes missing genotypes.

Pandora supports two distinct input data representations: file-based and *NumPy*based. For the file-based input, Pandora expects three distinct files per dataset: one genotype file containing the sequence data, one file containing metadata for each SNP, as well as one file containing metadata for each individual. Pandora supports the three file formats defined by the EIGENSOFT software package [17] (*EIGENSTRAT, ANCESTRYMAP, PACKEDANCESTRYMAP*), as well as the *pedigree*, and *binary pedigree* formats defined by the PLINK software package [20]. The *NumPy*-based representation relies on data matrices as defined by the Python library *NumPy* [9]. This allows users to preprocess their dataset using custom (Python) scripts without requiring an explicit (and potentially complicated as well as error-prone) conversion into a specific population genetics file format. Additionally, this input format allows for a more generic deployment of Pandora in other fields of biology beyond population genetics, such as PCA analyses of gene expression data, for instance.

Note that Pandora requires a preprocessed dataset, as it does not internally handle preprocessing. Thus, preprocessing steps, such as pruning Linkage Disequilibrium (LD) or rare variant filtering, must be completed *prior* to conducting Pandora analyses. This prerequisite does not constitute a limitation of Pandora as it applies to any Principal Component Analysis (PCA) or Multidimensional Scaling (MDS) analysis. For instance, EIGENSOFT’s *smartpca* tool also expects a preprocessed dataset when performing a PCA analysis. Thus, regardless of the specific analytical tool employed, conducting proper dataset preprocessing constitutes a fundamental step in population genetics research.

### 2.2 PCA and MDS

PCA and MDS are statistical methods to reduce the dimensionality in data that exhibit a high number of observations/features per individual. The goal is to reduce the dimensionality while preserving as much information as possible, in order to facilitate the interpretability and visualization of high-dimensional input data. While PCA directly operates on the genotype data, MDS reduces the dimensionality of a pairwise distance matrix. This distance matrix can be computed either on a per-individual or per-population basis. A commonly used distance metric is the genetic *F*_*ST*_ distance between populations [25], which reflects the proportion of total heterozygosity that can be explained by the within-population heterozygosity.

If the user provides the genotype data using the file-based representation, Pandora computes all PCA embeddings via EIGENSOFT’s *smartpca* tool, as it has been specifically designed and highly optimized for population genetics data [17, 19]. Therefore, Pandora supports and propagates all additional *smartpca* PCA analysis commands, including the option to project (ancient) individuals onto an embedding computed using another, distinct set of individuals. For MDS analyses, Pandora computes the *F*_*ST*_ distance matrix using *smartpca* and subsequently applies metric MDS as implemented in *scikit-allel* [15].

Using the alternative *NumPy*-based data representation, Pandora performs all PCA analyses using the PCA implementation provided in the Python machine learning library *scikit-learn* [18]. In analogy to the file-based data representation, the default distance metric is the *F*_*ST*_ distance (in this case as implemented in *scikit-allel*). Pandora also provides alternative distance metrics such as the Euclidean, Manhattan, or Hamming distances. The subsequent MDS is computed using the *scikit-allel* implementation of metric MDS.

Note that choosing the number of components for PCA or MDS constitutes a challenging and debated question. Patterson et al. [17] propose to base this decision on the Tracy-Widom statistic, but Elhaik [5] claims that this statistic is sensitive and results in overestimating the appropriate number of PCs. A common criterion used in other PCA application domains is to compute as many PCs as needed to explain, for instance, 80% of variance. However, this is not applicable to genotype data due to its extremely high dimensionality. For example, to explain 80% of the variance for a dataset analyzed by Lazaridis et al. [12], over 500 PCs need to be computed. In analogy to Price et al. [19], we thus set the default number of components for PCA analyses in Pandora to 10. For MDS analyses, we set the default number of components to 2. Pandora users can specify any number of components that best suit their analysis. Pandora uses all computed PCA or MDS components for its stability analyses. Further, note that the user needs to make an informed decision on the number of components in order to perform dimensionality reduction regardless of using Pandora.

To keep the following description of Pandora as generic as possible, we henceforth refer to the result of a PCA or MDS analysis as *embedding*.

### 2.3 Bootstrapping Genotype Data

The bootstrap procedure [4] is a statistical approach to quantify measures of accuracy based on random sampling with replacement from a set of observations. Estimating the stability of PCA using bootstrapping is not a novel concept. For example, Fisher et al. [7] propose a bootstrap-based method to estimate the sampling variability of PCA results on datasets where the number of per-sample measurements is substantially larger than the than number of samples. However, standard bootstrapping methods are not directly applicable to genotype data. A necessary condition for bootstrapping is the (assumed) independence of the bootstrapped measurements. Individuals (especially individuals of the same species) do, however, generally not evolve independently of each other. However, Felsenstein [6] argues that we may assume independent evolution of genetic loci. Therefore, in Pandora, instead of sampling the individuals, we sample the SNPs *s*_*i*_ to obtain a new bootstrap replicate matrix *B* ∈ ℝ^*M ×K*^. Thus, a bootstrap replicate has the exact same set of individuals but a distinct SNP composition. In doing so, we follow the standard approach used in phylogenetics, as first proposed by Felsenstein [6].

The computations of embeddings for individual bootstrap replicates are independent of each other, and their computations are therefore straight-forward to parallelize. Pandora exploits this parallelism and allows users to specify the number of threads to use.

Since Pandora bootstraps an embedding (a compute-intensive dimensionality reduction), the computational resource requirements are substantial. To alleviate this, we periodically perform a heuristic convergence assessment until a maximum number of bootstrap replicates is reached. The frequency of this convergence check is determined by the number of threads being used. During the convergence check, Pandora assesses the variation of PS estimates based on random subsets of the computed replicates at the time of the convergence check. When this variation is below a certain tolerance level, Pandora terminates all remaining bootstrap computations and conducts the final stability value calculation using the set of completed bootstrap replicates. We provide a detailed description of our convergence criterion in Supplementary Material Section 1.1. The default maximum number of bootstrap replicates in Pandora is 100 and the default convergence tolerance is 0.05. Both settings can be adjusted by the user.

### 2.4 Stability Estimation

#### 2.4.1 Pandora Stability

The Pandora Stability (PS) describes the overall stability of the genotype dataset that is based on the pairwise similarity scores over all bootstrap replicates. To compute the similarity between two bootstrap embeddings *P*^*u*^ and *P*^*v*^, we follow the approach suggested by Wang et al. [24] using Procrustes Analysis. Procrustes Analysis determines the optimal transformation *f* (*P*^*u*^) that matches the projections of all individuals *x*_*i*_ in *P*^*u*^ to *P*^*v*^ as accurately as possible while preserving the relative distances between individuals. This transformation consists of a scaling factor, a rotation and reflection matrix, and a translation matrix. In Pandora, we use the Procrustes Analysis as implemented in the Python scientific computing package *SciPy* [23]. Prior to computing the optimal transformation, both embeddings are standardized such that *trace*(*PP*^*T*^) = 1, and both sets of points are centered around the origin to remove translation effects. The optimal transformation minimizes the squared Frobenius norm (also called disparity)

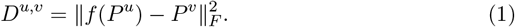

The disparity *D*^*u,v*^ has a minimum of 0 and a maximum of 1 [24]. Based on this *disparity*, we can compute the pairwise *similarity* as

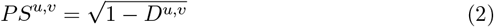

on a scale from 0 to 1, with higher values indicating higher similarity [24]. To obtain the overall PS, we average over the pairwise similarity across all 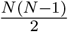 possible distinct pairs of bootstrap replicates

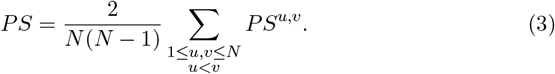

The resulting PS is again a value between 0 and 1. The higher the PS, the more similar the bootstrap replicates are, with 1 indicating that all bootstrap embeddings identically project all individuals.

#### 2.4.2 Pandora Cluster Stability

In analogy to the PS, we initially compute the pairwise Pandora Cluster Stability (PCS) for all unique pairs of bootstrap embeddings and then average over all pairwise scores to obtain the overall PCS

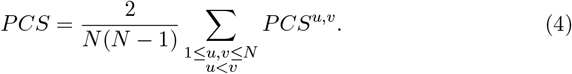

To compute the pairwise *PCS*^*u,v*^ for bootstrap embeddings *P*^*u*^ and *P*^*v*^, we first match both embeddings using Procrustes Analysis as described above and subsequently perform an independent *k*-means clustering for the two embeddings. In order to compare the resulting label assignments for all individuals, we use the same number of clusters *k* for both embeddings. Finally, we compute the pairwise *PCS*^*u,v*^ for bootstrap embeddings *P*^*u*^ and *P*^*v*^ via the Fowlkes-Mallows index [8].

The PCS is a value between 0 and 1. The higher the PCS, the higher is the similarity between clustering results across bootstraps.

Selecting the number of clusters *k* to best represent the dataset is not trivial. Pandora users can manually set the number of clusters based on, for instance, prior knowledge of the expected underlying (sub-)population structure in the data. When the user does not explicitly specify *k*, we automatically determine the optimal *k* based on a grid search and the Bayesian Information Criterion (BIC) [21]. To reduce the computational cost of searching for the optimal *k*, we determine an input data specific upper bound via the following heuristic. If the individuals of the dataset to be analyzed have population annotations, we set the upper bound to the number of distinct populations in the dataset. If this information is not available, we set the upper bound to the square root of the number of individuals in the dataset. Based on empirical parameter exploration experiments, we set the lower bound to 3. During Pandora development, we observe a substantial influence of *k* on the resulting PCS scores for various empirical population genetics datasets. We provide further details on these experiments in Supplementary Material Section 2.4.

#### 2.4.3 Pandora Support Values

To assess the projection stability for each individual in the dataset, we compute the Pandora support for each individual *x*_*i*_ as

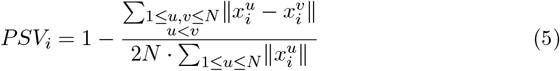

where 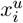 and 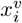 denote the projection of *x*_*i*_ in bootstrap *u* and *v* respectively.

The fraction in Equation (5) corresponds to the Gini coefficient, a statistical measure of dispersion from economics. In our case, the Gini coefficient measures the dispersion of the projections of an individual with respect to all bootstrap replicates on a scale from 0 to 1. The higher the dispersion, the higher the Gini coefficient will be. To align this with the interpretation of the PS and PCS values, where 0 corresponds to unstable and 1 to stable, we simply subtract the calculated dispersion measure from 1.

## 3 Discussion

In the following, we demonstrate the utility of Pandora. Due to the novelty of our approach, and the novelty of stability estimation of genotype data under dimensionality reduction, there exists no benchmark data collection to compare Pandora against. We therefore verify and demonstrate the functionality of our proposed approach using simulated genotype data and empirical, published population genetics datasets. Note that we provide all scripts to reproduce the following analyses on GitHub https://github.com/tschuelia/PandoraPaper and all results can be downloaded at https://cme.h-its.org/exelixis/material/Pandora_supplementary_data.tar.gz. We additionally provide further information on the software versions and hardware we used in Supplementary Material Section 3.

### 3.1 Simulated Data

We first analyze the functionality of bootstrap-based stability estimation using simulated genotype datasets. We simulated these datasets using the *stdpopsim* python library [1, 11]. This library provides a catalog of 13 distinct, published human demographic models describing the demographic history of various human populations. We simulated genotype data according to each demographic model to generate realistic whole-genome population genetic data. See Supplementary Material Section 2.1 for a detailed description of our simulation and analysis setup.

For the following analyses, we estimated the stability using Pandora with the default bootstrap convergence setting (convergence tolerance of 5%). For PCA analyses, we set the number of components to 10, and we disabled outlier detection in *smartpca*. For MDS analyses, we reduced the data to 2 dimensions, and we used the Euclidean distance between individuals as the distance metric.

For our first analysis, we simulated data with varying sequence lengths (10^5^, …, 10^8^). Simulating longer genomes increases the number of SNPs in the simulated dataset. With increasing SNP numbers, we expect the amount of informative data in the genotype dataset to also increase. Thus, we expect dimensionality reduction to better capture population structure. Consequently, we expect PCA or MDS results to become more stable with increasing SNP numbers in the dataset. Note that while we simulate sequences of length 10^5^, this will not necessarily yield 10^5^ SNPs since not all genomic sites evolve. Supplementary Table S1 summarizes the number of SNPs per simulated sequence length for all simulated datasets. Since the resulting SNP number varies as a function of the demographic model, in the following, we identify datasets via the *target* sequence length. Figure 2 shows the distribution of PS values for PCA analyses as a function of the target sequence length (see Supplementary Figure S5 for the respective results of the MDS analyses). The expected trend of higher stability with increasing target sequence lengths is clearly visible for both, PCA, and MDS. This trend can be visualized by plotting the first two principal components of the PCA analysis per target sequence length. Supplementary Figure S2 shows this trend for the demographic model of ancient European populations.

**Figure 2:**
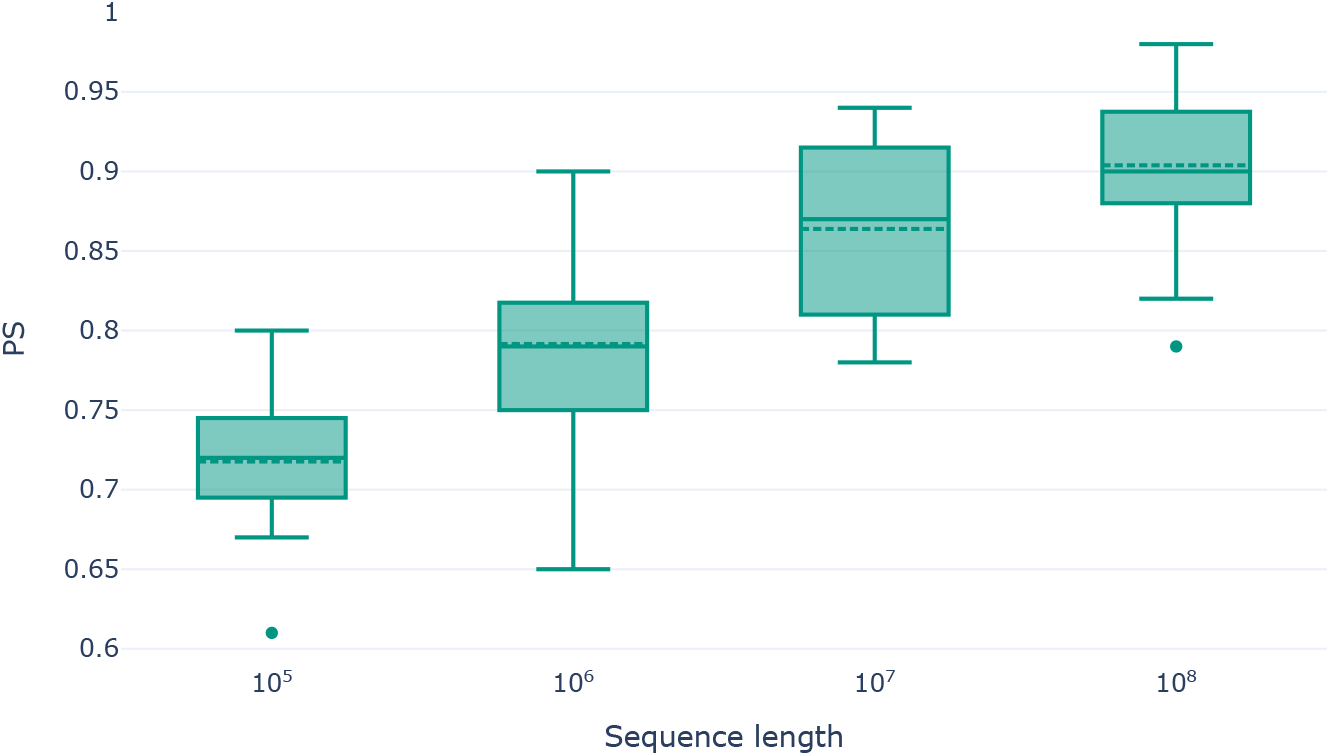
PS values for PCA analyses as a function of the target sequence length for the simulated genotype datasets.

Missing data is known to affect the stability of dimensionality reduction [26]. We therefore expect decreasing PS values with increasing proportions of missing data. To test this hypothesis, we distorted simulated datasets by injecting missing data into our datasets with a target sequence length 10^8^. Note that we used our datasets with the maximum target sequence length to not bias the following analysis by low(er) signal resulting from small SNP numbers in the dataset. For the missing data impact analysis, we altered the dataset by replacing proportions of the data (1%, 5%, 10%, 20%, and 50%) with the missing data character. Note that this does not alter the number of SNPs per dataset. Further, note that imputing (or removing) missing data is essential for PCA or MDS analyses, as the underlying mathematical frameworks cannot process incomplete datasets. Thus, we imputed missing data using mean imputation.

Figure 3 shows the distribution of PS values as a function of the missing data proportion. We do observe the expected trend of decreasing PS values with increasing missingness. Yet, even with 50% missing data, the results are relatively stable with an average PS value of 0.78, and its decline is not as substantial as one might expect. One reason for this is that the simulated datasets are overall clearly structured with highly distinct populations. For example, Supplementary Figure S3 shows the first two principal components of the dataset simulated according to the demographic model of ancient European populations with increasing proportions of missing data. Despite the high proportion of missing data of 50% (bottom right panel), the population structure was easily recovered. We observe only a slight dispersion within the individual population clusters compared to the PCA on the dataset without missing data (upper left panel). Furthermore, Yi and Latch [26] report analogous findings: while PCA results are affected by random missing data, high proportions of random missing data can be compensated by a strong underlying population structure.

**Figure 3:**
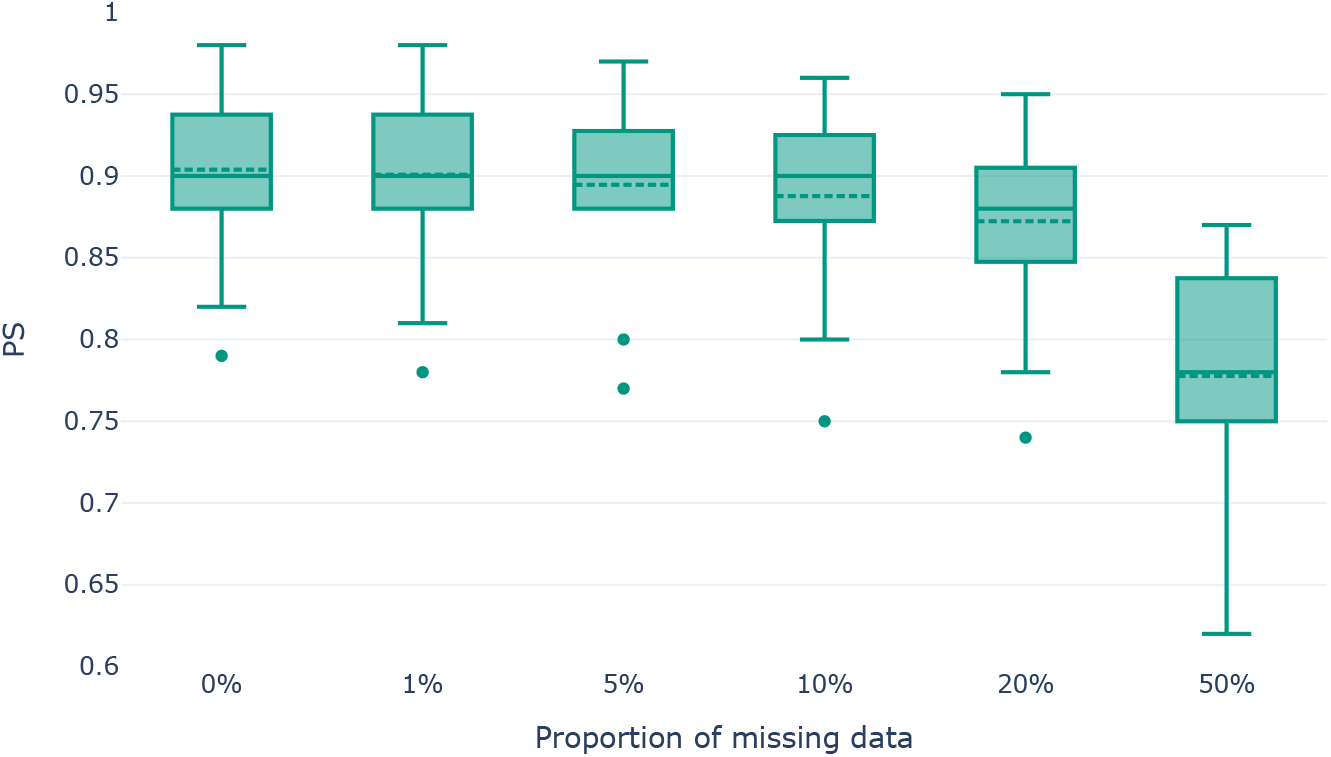
PS values for PCA analyses as a function of the proportion of random missing data for the simulated genotype datasets.

For MDS analyses, we observe no substantial effect of missing data on the stability of genotype datasets for missing data proportions up to 20%. Even for 50% missing data, there is only a slight PS decrease (see Supplementary Figure S6). Evidently, the distance matrix calculation is fairly robust to the imputation of missing data, as is the MDS embedding construction with respect to small changes in the distance matrix.

In contrast to *missing* data, *noisy* data cannot be imputed, as we do not know which part of the data is noise and which part is “real” data. We expect lower PS values for higher proportions of noise in the dataset because uninformative noise distorts the structure of a dataset. For the following simulations, we used our simulated datasets with target sequence length 10^8^ and without any missing data. We distorted the data by adding 10%, 20%, and 50% of noise. To this end, we altered single genotypes of randomly selected SNPs and individuals in the genotype matrix.

As expected, we observe a substantial decline in stability with higher proportions of random noise. This observation holds true for both PCA (see Figure 4) and MDS (see Supplementary Figure S7) analyses. However, the effect is substantially more pronounced in PCA analyses. Using the simulated dataset of the demographic model of ancient European individuals, Supplementary Figure S4 visualizes the impact of noise on the resulting population structure in the PCA, clearly showing a higher dispersion of individuals.

**Figure 4:**
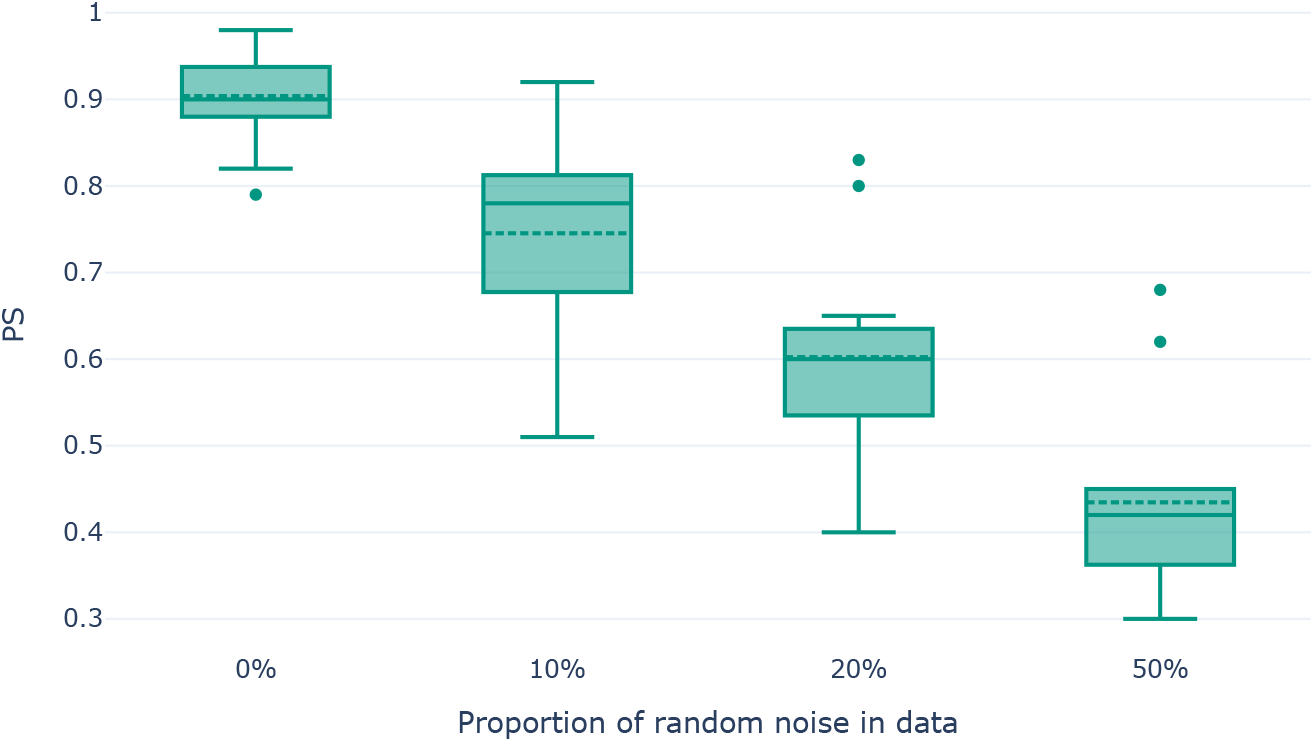
PS values for PCA analyses as a function of the proportion of random noise data for the simulated genotype datasets.

Given that Pandora consistently reflects the expected trends of increasing stability with an increasing number of SNPs, and decreasing stability with increasing proportions of missing and noisy data, we conclude that bootstrapping-based stability estimation properly captures instability in dimensionality reduction studies.

### 3.2 Empirical Data

All the simulated datasets exhibit a clear underlying population structure. As this is not necessarily the case for empirical data, we also demonstrate the utility of Pandora on empirical datasets. Additionally, PCA or MDS studies are frequently applied in ancient DNA (aDNA) studies. Usually, in aDNA studies, the embedding is computed using a dataset of modern individuals. Subsequently, ancient individuals are projected onto the resulting PCA or MDS embedding. In human population studies, especially in work on newly sequenced ancient specimen, the Human Origins (HO) SNP Array [12] comprising 2068 publicly available individuals is used as reference dataset of modern individuals. For our first empirical analysis, we assembled the West-Eurasian subset based on the description provided by Lazaridis et al. [12]. The resulting West-Eurasian subset contains 813 individuals from 55 distinct populations. We further denote this dataset as *HO-WE*. The *HO-WE* dataset contains 605 775 SNPs with 0.5% missing data. We then merged the *HO-WE* dataset with the 13 ancient Çayönü individuals published by Altınışık et al. [2] (*HO-WE-Çayönü*). These ancient individuals contain between 90% and 99% missing data. Using the *smartpca* option `lsqproject:YES`, we computed all PCAs in the following analysis using the modern individuals only and then projected the 13 ancient Çayönü individuals onto the resulting embeddings. According to Pandora, the *HO-WE-Çayönü* dataset is overall very stable under PCA, with a PS of 0.89. In their analyses, Altınışık et al. [2] computed a 95% confidence region for the first two principal components using the confidence ellipse function in *smartpca*. This confidence interval indicates a high projection uncertainty, especially for the *cay015* individual. Our results are in line with these findings, as the *cay015* individual has a PSV of only 0.54 according to our analyses. Figure 5 visualizes this behavior, showing the computed PCA embeddings of two bootstrap replicate datasets and highlighting the *cay015* individual. A comparison of the figures clearly shows that the positions of the *cay015* individual in both embeddings differ substantially, hence the low PSV. For individuals with such low PSVs, conclusions based on PCA analyses should only be drawn with extreme caution. For example, *cay015* is projected closest to an Israeli individual in the PCA of the unbootstrapped *HO-WE-Çayönü* dataset (*BedouinA* population). However, when considering the two bootstraps we used in Figure 5, *cay015* is projected closest to a Yemeni individual (bootstrap replicate #1; *Yemenite-Jew* population) or a Libyan individual (bootstrap replicate #2; *Libyan-Jew* population). These observations are again in line with the findings of Altınışık et al. [2], as they conclude that Çayönü is a genetically diverse population.

**Figure 5:**
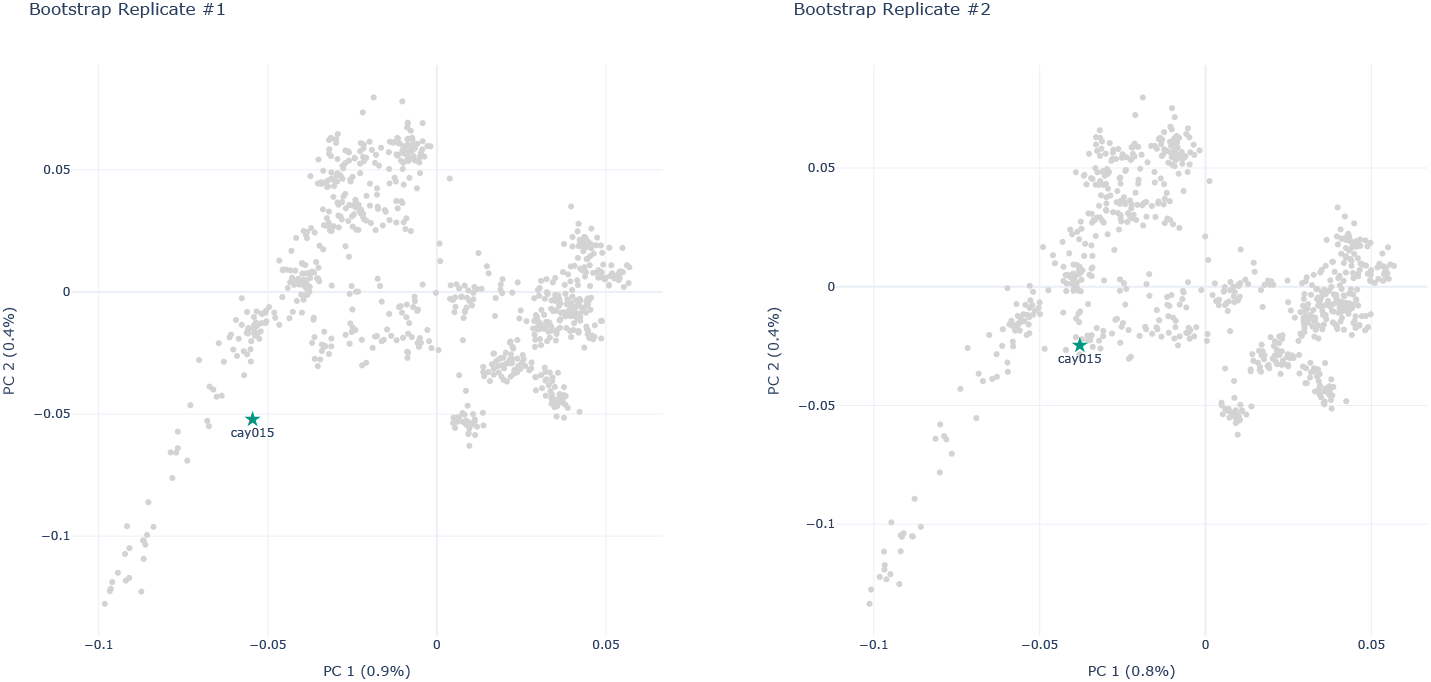
PCA embeddings of two bootstrap replicates of the *HO-WE-Çayönü* dataset The highlighted points correspond to the respective projection of the *cay015* individual.

We suggest filtering individuals with low PSVs prior to any additional analyses. If, however, the individuals with low PSVs are important to the study, we suggest performing additional analyses, for instance admixture analyses. Since version 8, *smartpca* includes a confidence ellipse functionality that plots a 95% confidence region for individuals. This functionality can be especially useful for the placement of ancient individuals (see for example the analyses in Altınışık et al. [2]). However, the confidence ellipse only considers the first two principal components and does not *quantify* the instability, but yields a plot showing the confidence regions. Consequently, this approach cannot be performed simultaneously for all individuals in the dataset, as the resulting figure would be overly complicated. With Pandora, users can pre-filter instable individuals with low PSVs and subsequently perform a more targeted confidence region analysis.

Finally, we analyzed two datasets of genomic dog data published by Morrill et al. [16]. Using the respective definition of Morrill et al. [16], we assembled a subset of 601 *purebred* dogs and a subset of 1071 *highly admixed* dogs based on the 2155 publically available genomic sequences comprising 6 059 222 SNPs. The idea of this analysis was to demonstrate the usage and utility of Pandora’s PCS estimate. We determined the number of distinct breeds in each dataset using the data provided by Morrill et al. [16]. The purebred dogs comprise 88 distinct breeds, whereas the highly admixed dogs comprise 60 distinct dog breeds. We assessed the stability of breed assignments via the PCS as estimated by Pandora, using the respective number of breeds as *k* in *k*-means clustering. Morrill et al. [16] consider a dog as being *highly admixed* if it has below 45% ancestry from any single breed in admixture analysis. Consequently, we expect a lower PCS for the *highly admixed* dataset compared to the *purebred* dataset. In fact, for the *highly admixed* dataset we observe a PCS of only 0.63, whereas the *purebred* dataset is substantially more stable under *k*-means clustering with a PCS of 0.87.

### 3.3 Speedup and Accuracy under the Bootstrap Convergence Criterion

Using the 13 simulated population genetics datasets (simulated with target sequence length 10^8^ and no missing or noisy data), we assessed the impact of the bootstrap convergence criterion on the runtime and accuracy of Pandora. We compared the runtime and deviation of PS values, as well as PSVs when performing Pandora analysis using the default 5% convergence tolerance setting and a lower 1% convergence tolerance setting to the respective Pandora execution with convergence checks disabled and computing the full set of 100 replicates (see Supplementary Material Section 2.3 for further details).

For PCA analyses, we observe an average speedup of 2.6 ± 0.6 for the 5% convergence tolerance. Bootstrap convergence occurred after computing 20 replicates for all 13 datasets. With the 1% convergence tolerance, we observe an average speedup of 1.4 ± 0.4 after 63 replicates on average. The PS values deviate only slightly, with a maximum observed deviation of 0.01 for both settings. The PSVs of individuals differ on average by 0.01 with the 5% convergence tolerance and by 0.0 with the 1% convergence tolerance. We observe a maximum deviation of PSVs for single individuals of 0.08 and 0.06 respectively.

The speedup and deviation of PS values are similar for MDS analyses. We observe an average speedup of 2.2 ± 0.6 with the 5% convergence tolerance, and an average speedup of 1.7 ± 0.7 with the 1% convergence tolerance. Pandora converged on average after 20 (5%) and 34 replicates (1%). In analogy to the PCA results, the PS deviations for MDS deviate only slightly, with maximum deviations of 0.02 for both convergence tolerance settings. The average PSV deviation is 0.01 under both settings. However, we observe substantial deviations for single individuals, with differences up to 0.19.

Our results suggest that for PCA and MDS analyses, the 5% convergence tolerance is suitable when the overall PS is of primary interest. When correct PSVs estimates are paramount, we recommend decreasing the convergence tolerance to 1% for PCA analyses, and disabling the convergence check for MDS analyses.

See Supplementary Material Section 2.3 for box plots of the speedup, PS, and PSV deviations summarized for all 13 datasets.

### 3.4 Summary

Pandora estimates the stability of a genotype dataset and its individuals under dimensionality reduction via bootstrapping. Pandora supports PCA and MDS analyses. In addition to an overall stability estimate (PS), Pandora applies *k*-means clustering to all bootstrapped embeddings to estimate the stability of the assigned clusters across all bootstrap replicates (PCS). Furthermore, Pandora computes a bootstrap support value for each individual in the genotype dataset (PSVs). The latter is particularly useful for datasets with projections of ancient individuals onto an embedding computed on modern individuals only. For these analyses, the PSVs for the ancient individuals provide a confidence value regarding their placement in the embedding space relative to modern individuals. The lower the support for an individual, the more careful users should be about drawing conclusions regarding this individual based on dimensionality reduction analyses.

Our analysis results on empirical and simulated data show that Pandora is able to detect instability in PCA and MDS analyses. As expected, Pandora yields lower stability for both, PCA and MDS analyses, with increasing fractions of missing and noisy data under simulations, as well as higher stability with increasing number of SNPs in the dataset. Compared to a single PCA or MDS analysis, Pandora introduces a substantial runtime and resource overhead. Our bootstrap convergence criterion alleviates this to the extent possible.

Our analyses using simulated genotype datasets showed that Pandora is particularly useful when performing PCA or MDS analyses for datasets with few SNPs, high proportions of missing data, or if the data is expected to be noisy. With the exemplary analysis of the empirical *HO-WE-Çayönü* dataset, we further demonstrated the utility of Pandora stability values for preventing potentially erroneous and hence misleading conclusions based on unstable dimensionality reduction.

We believe that Pandora will be of high value to population genetic studies as the inherent uncertainty of PCA and MDS analyses can now be seamlessly and routinely assessed via an easy-to-use tool and thereby contribute to circumvent potentially biased conclusions.

In Pandora, the bootstrapping and the subsequent dimensionality reduction are both performed along the SNPs. Pandora does not implement a resampling of the individuals, since Elhaik [5] demonstrated the sensitivity of PCA studies to (subjective) selection of individuals. The author showed that PCA analyses can be manipulated to confirm the desired hypothesis, depending on the selection of individuals used for PCA analyses. Consequently, for Pandora, we refrained from sampling individuals. Thus, Pandora results can not directly be used to justify a subjective selection of individuals. However, such an individual-sampling-based procedure might allow estimating the bias induced by specific individuals. In conjunction with the Pandora stability estimates, especially the PSVs, this can contribute to identify an objective, meaningful set of individuals for population genetics studies. Devising such a selection criterion will constitute the subject of future work.

As previously mentioned, despite their popularity, the use of PCA or MDS in population genetics is controversial. More modern dimensionality reduction approaches such as t-SNE [22], UMAP [13], or deep learning based variational autoencoders [3] alleviate some problems, but exhibit inherent disadvantages (see Battey et al. [3] for a discussion). Integrating such methods into Pandora as alternatives to PCA or MDS also constitutes future work.

## 4 Acknowledgements

We thank N. Ezgi Altınışık for the very helpful discussion on Pandora, as well as the valuable insights in the methodology of population genetics studies. We thank the aDNA lab at IMBB-FORTH in Heraklion, Crete for the useful initial discussions and ideas. We further thank Jonas Haag for the invaluable help in debugging Python multiprocessing. This work was financially supported by the Klaus Tschira Foundation, by the European Union (EU) under Grant Agreement No 101087081 (Comp-Biodiv-GR), and by the Helmholtz Pilot Program Core-Informatics at KIT (KiKIT).

