## Supplementary Material for "Pandora: A Tool to Estimate Dimensionality Reduction Stability of Genotype Data"

February 2025

### 1 Extended Methods

#### 1.1 Bootstrap Convergence Criterion

Since Pandora bootstraps an embedding (a compute-intensive dimensionality reduction), the resource usage and compute time is substantial. To reduce this computational overhead, we periodically perform a heuristic convergence assessment until a maximum number of bootstrap replicates is reached. The frequency of this convergence check is determined by the number of threads used.

The default setting is a maximum of 100 bootstrap replicates. Now, let $N^*$ be the number of bootstrap replicates already computed when performing the convergence assessment. We create 10 subsets of size $\tilde{N} = \frac{N^*}{2}$ by sampling 10 times without replacement from the set of $N^*$ completed bootstraps. We then compute the Pandora Stability (PS) for each of the 10 subsets and determine the relative difference of PS values between all pairs of PS values $(PS_u, PS_v)$: $\frac{|PS_u - PS_v|}{PS_u}$. We assume convergence if no pairwise relative difference exceeds $\alpha$. In other words, we require that the highest sampled stability is at most $\alpha * 100\%$ larger than the lowest sampled stability. If this is the case, all remaining

bootstrap computations are omitted. Pandora allows users to explicitly set the convergence tolerance $\alpha$; the default setting is 0.05. Note that we decided to sample 10 subsets of size $\tilde{N}$ as a trade-off between the accuracy and the additional runtime overhead induced by the bootstrap convergence assessment. The runtime overhead induced by the convergence assessment is substantial, as we need to compute the PS for each of the subsets, meaning that we need to perform $\frac{\tilde{N}(\tilde{N}-1)}{2}$ Procrustes Transformations.
We also determine the frequency of this convergence assessment based on the number of threads specified by the user, which defines the size of a full batch of bootstrap replicates. Pandora checks for convergence after every full batch or once 10 replicates are computed, whichever number is higher. For example, if a user executes Pandora with 40 threads, it will compute 40 bootstrap replicates in parallel and hence complete 40 replicates at approximately the same time. If Pandora were to run the convergence assessment after completing 10 bootstraps and determined that it has converged, we would discard 30 (almost) finished bootstraps, thus wasting the resources used for their computation, as well as for three unnecessary convergence assessment computations (after 10, 20, and 30 completed replicates).

### 2 Extended Discussion

#### 2.1 Data Simulation

We simulated genotype datasets using the *stdpopsim* python library [1, 4]. This library provides a catalog of 13 distinct, published human demographic models describing the demographic history of various human populations. We simulated genotype data according to each model to generate realistic whole-genome population genetic data. Note that we set *msprime* [2] as simulation engine for all following *stdpopsim* simulations. To not bias the simulations in favor of single populations, we simulated the same number of individuals for each population in the demographic model, up to a total of 500 individuals.

Figure S1 visualizes our simulation pipeline. We first simulated genotype datasets with a *target* sequence length of $10^5$, $10^6$, $10^7$, and $10^8$ nucleotides. While the resulting simulated sequences are of this target length, not all positions in the simulated sequences are SNPs. Consequently, we only used the SNPs for the subsequent analyses. We filtered variants with an allele frequency below 1% (*MAF filtering*). Additionally, we removed correlated SNPs by applying Linkage Disequilibrium (LD) pruning with a $r^2$ threshold of 0.5, a window size of 50 and a stride of 5. We performed LD pruning and MAF filtering using PLINK 2.0 [3]. Table S1 states the resulting number of SNPs per simulated dataset (by model and target sequence length).

Using our 13 datasets simulated with a target sequence length of $10^8$, we distorted the data using random missing data and random noise. For the missing data simulations, we randomly replaced a certain proportion in the genotype matrix with the missing value character (1%, 5%, 10%, 20%, and 50%). For

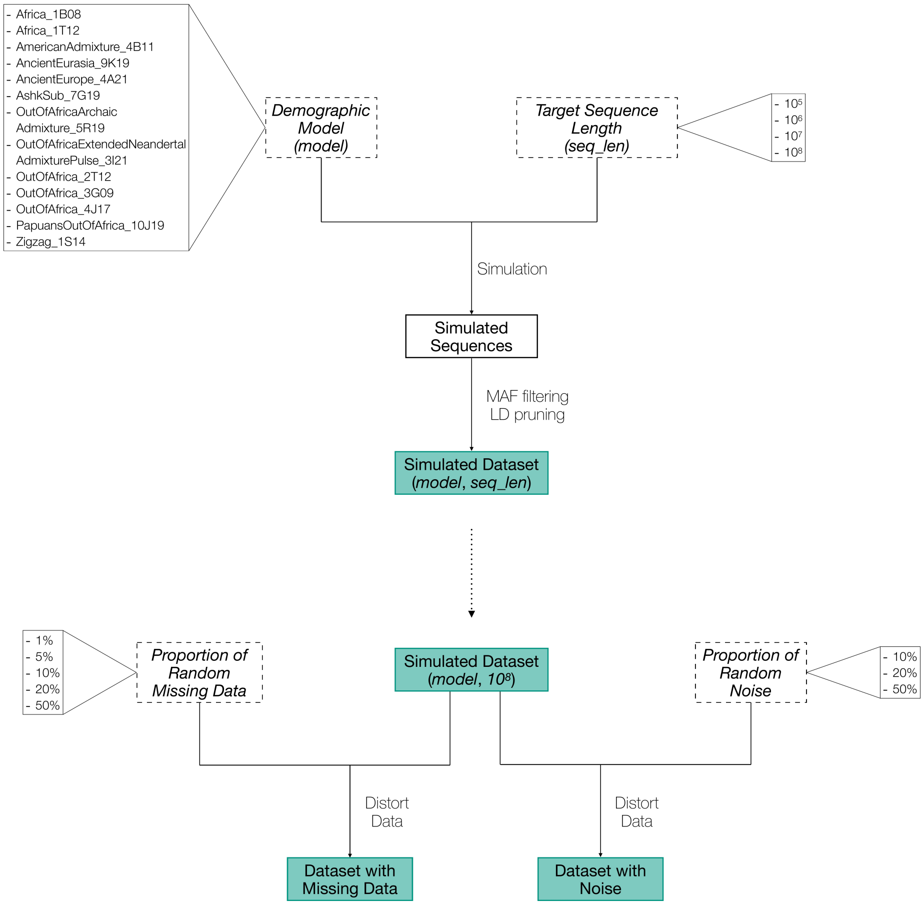


Figure S1: Schematic overview of the implemented simulation pipeline.

the simulations of noisy data, we randomly replaced a certain proportion of values in the genotype matrix with random variants (10%, 20%, and 50%). The number of SNPs remains unaffected by these changes. Note that we did not mix missing and noisy data simulations. That is, for the missing data simulations we did not add additional noise, and for the noise data simulation we did not add missing data.

We performed Pandora stability analyses using PCA and MDS for all datasets highlighted in Figure S1. For PCA analyses, we reduced the data to 10 dimensions (the default setting in *smartpca*). For MDS analyses, we reduced the data to 2 dimension, and we used the Euclidean sample distance and distance metric. We used 20 threads for all Pandora analyses.

We implemented the outlined simulation pipeline using the *Snakemake* workflow management system [5]. Note that we set the random seed for each simulation to ensure reproducible simulation results. We provide all scripts required

| Demographic Model | Target Sequence Length | | | |
|---|---|---|---|---|
| | $10^5$ | $10^6$ | $10^7$ | $10^8$ |
| Africa_1B08 | 217 | 2255 | 22 172 | 222 290 |
| Africa_1T12 | 195 | 1924 | 18 924 | 190 410 |
| AmericanAdmixture_4B11 | 165 | 1470 | 13 076 | 135 622 |
| AncientEurasia_9K19 | 142 | 1331 | 13 446 | 133 883 |
| AncientEurope_4A21 | 60 | 692 | 6722 | 65 006 |
| AshkSub_7G19 | 183 | 2202 | 21 573 | 207 071 |
| OutOfAfrica-ArchaicAdmixture_5R19 | 105 | 1019 | 10 546 | 106 972 |
| OutOfAfricaExtended-NeandertalAdmixturePulse_3I21 | 272 | 2920 | 29 971 | 296 220 |
| OutOfAfrica_2T12 | 182 | 1769 | 16 721 | 163 492 |
| OutOfAfrica_3G09 | 153 | 1560 | 15 725 | 156 819 |
| OutOfAfrica_4J17 | 127 | 1502 | 14 477 | 142 285 |
| PapuansOutOfAfrica_10J19 | 411 | 3994 | 38 065 | 373 325 |
| Zigzag_1S14 | 150 | 1586 | 17 449 | 171 090 |

Table S1: Number of SNPs in the simulated population genetics datasets per target sequence length.

for reproducing our simulations on GitHub (`https://github.com/tschuelia/PandoraPaper`) and all simulated datasets (including all datasets with random missing and noise data) in EIGENSTRAT format at `https://cme.h-its.org/exelixis/material/Pandora_supplementary_data.tar.gz`.

#### 2.2 Simulated Data

##### 2.2.1 PCA Stability Analyses

Figure S2 visualizes the first two principal components of the PCA computed for data simulated under the AncientEurope_4A21 demographic model with increasing target sequence lengths ($10^5$, . . . , $10^8$). The longer the target sequence, the better the population structure becomes visible. As expected, the PS increases with increasing sequence lengths.

Figure S3 visualizes the first two principal components of the PCA computed for data simulated under the AncientEurope_4A21 demographic model with a target sequence length of $10^8$ and increasing proportions of random missing data (1%, 5%, 10%, 20%, and 50%). The PS is identical for up to 10% missing data. With 20% missing data, we observe a slight PS decrease (from 0.88 to 0.84) and for 50% missing data, the PS decreases further to 0.76. Yet, even with 50% missing data, PCA was able to easily detect the underlying population structure. As stated in the main paper, random missing data (especially when missing data is imputed in PCA analyses), does not have a substantial impact on the stability of PCA analyses if the underlying population structure is strong.

Figure S4 visualizes the first two principal components of the PCA computed

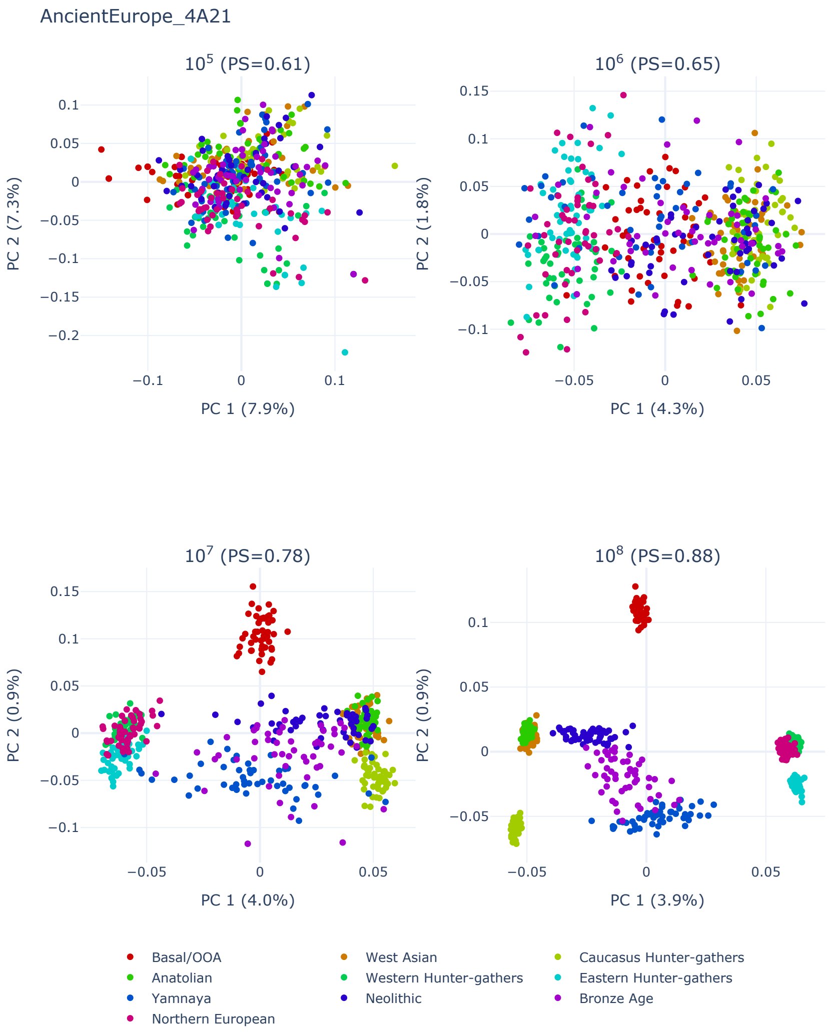


Figure S2: First two principal components of the PCA computed for data simulated under the AncientEurope_4A21 demographic model with increasing target sequence lengths.

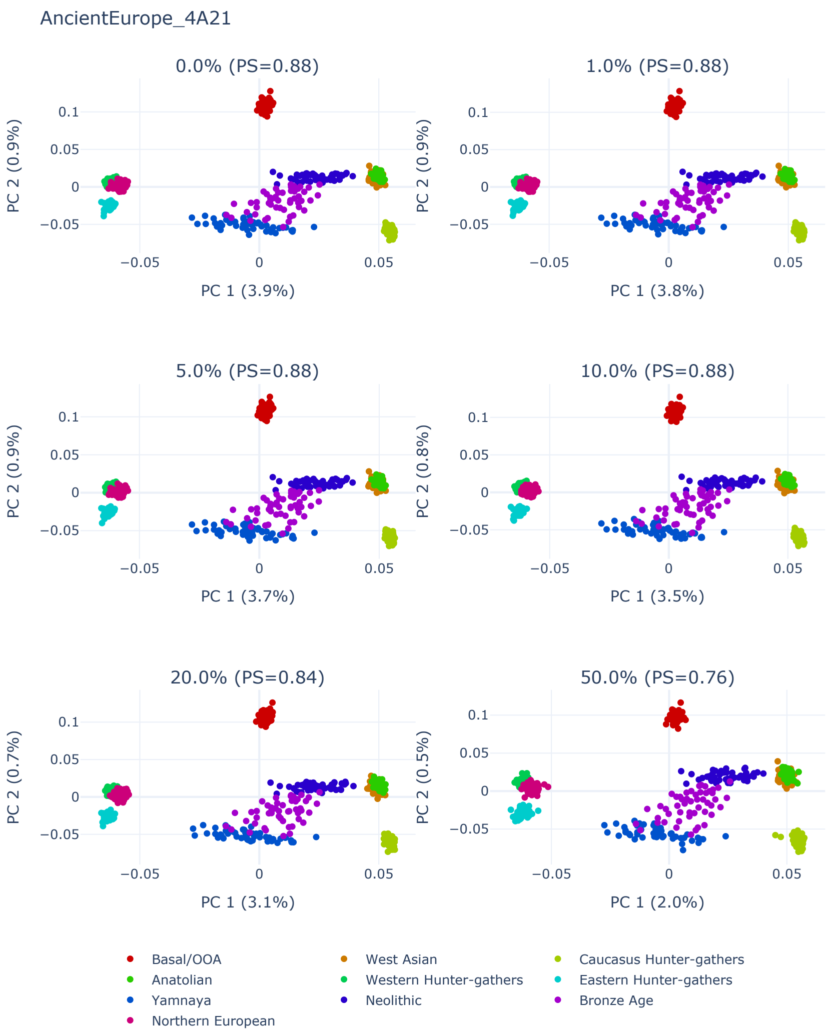


Figure S3: First two principal components of the PCA computed for data simulated under the AncientEurope_4A21 demographic model with a target sequence length of $10^8$ and increasing proportions of random missing data.

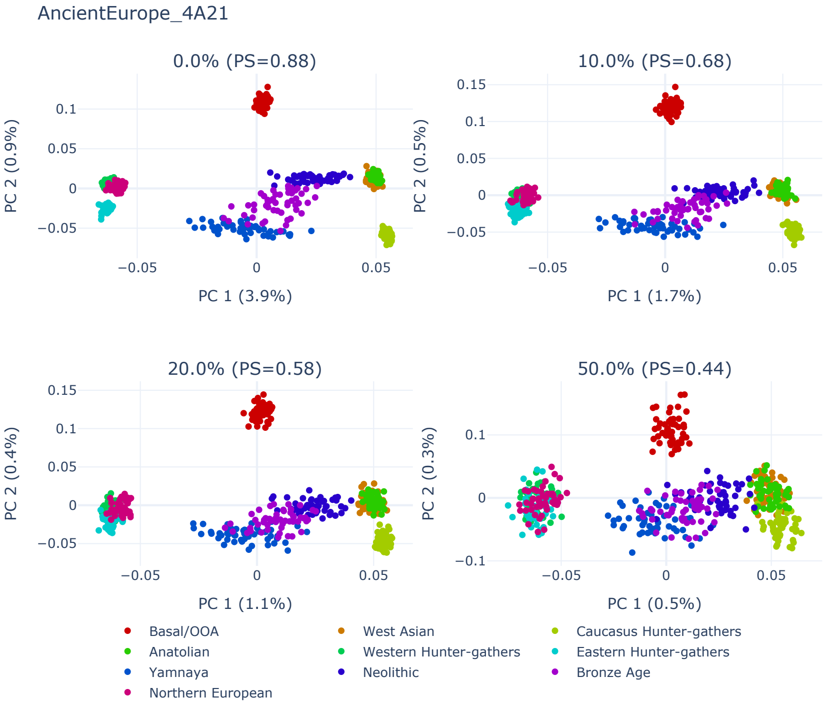


Figure S4: First two principal components of the PCA computed for data simulated under the AncientEurope_4A21 demographic model with a target sequence length of $10^8$ and increasing proportions of random noise.

for data simulated under the AncientEurope_4A21 demographic model with a target sequence length of $10^8$ and increasing proportions of random noise (10%, 20%, and 50%). As expected, the PS decreases with higher proportions of noise.

##### 2.2.2 MDS Stability Analyses

For all MDS analyses in this section, we reduced the data to 2 dimensions using MDS with the Euclidean sample distance as the distance metric. We estimated the stability using Pandora with the default convergence tolerance setting of 5%.

Figure S5 shows the PS values of the simulated datasets as a function of the target sequence length of the dataset simulation. In analogy to the PCA results, we observe an increased PS with longer target sequence lengths.

Figure S6 shows the PS values of the simulated datasets with increasing proportions of missing data.

Finally, we estimated the stability of MDS analyses as a function of the proportion of random noise in the data. Figure S7 shows that, similar to the

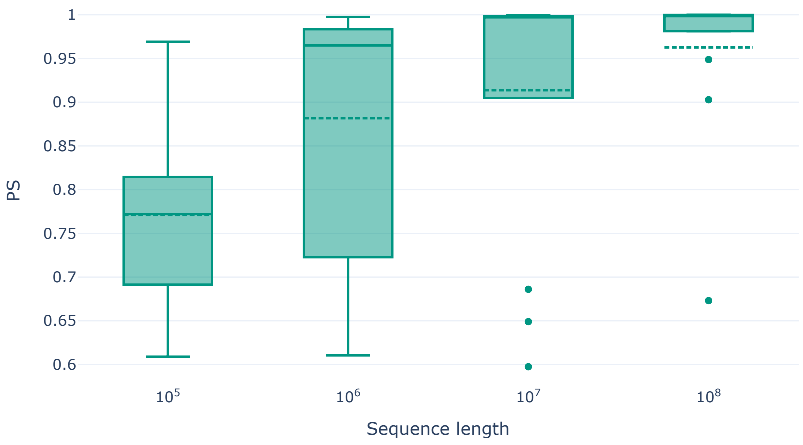


Figure S5: PS values for MDS analyses as a function of the target sequence length for the simulated genotype datasets.

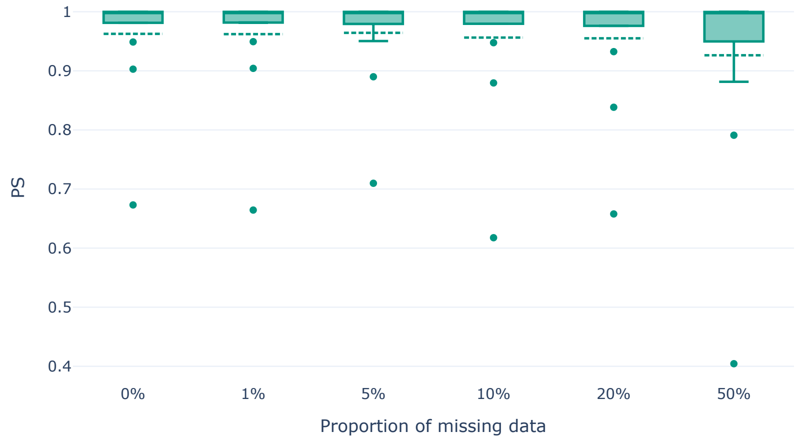


Figure S6: PS values for MDS analyses as a function of the proportion of random missing data for the simulated genotype datasets.

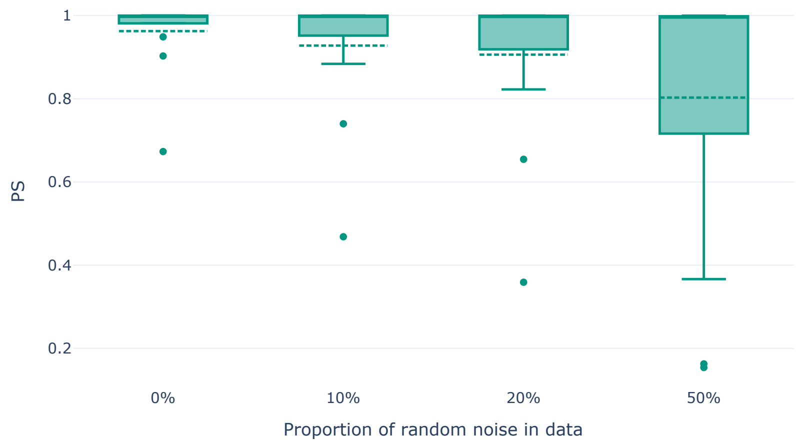


Figure S7: PS values for MDS analyses as a function of the proportion of random noise data for the simulated genotype datasets.

PCA analyses in the main paper, we observe a decrease of PS values with higher proportions of noise. However, the effect is substantially less pronounced than with PCA analyses. In analogy to the argument above, we suspect MDS to be less prone to distorted data due to the distance matrix based approach.

#### 2.3 Speedup and Accuracy under the Bootstrap Convergence Criterion

In the following, we describe our analysis setup to assess the influence of the implemented bootstrap convergence check as described in Section 1.1 on the stability estimates and the runtime of Pandora. For each of the 13 simulated genotype datasets (target sequence length $10^8$ and no missing or noisy data), we performed the following analyses. As baseline, we executed Pandora with the full set of 100 bootstrap replicates (default Pandora setting) and disabled the bootstrap convergence check. We denote the runtime of this execution by $T_{100}$, the PS as $PS_{100}$, and the PSVs as $PSV_{100}$. We further executed Pandora twice with the convergence check enabled, but with a different convergence tolerance in each execution. In one execution, we set the convergence tolerance to 0.05 (5%) and in a second execution to a more conservative value of 0.01 (1%). We denote the results of both executions as $T_{5\%}, PS_{5\%}, PSV_{5\%}$ and $T_{1\%}, PS_{1\%}, PSV_{1\%}$, respectively. We further report the number of bootstraps required for convergence for both executions ($N_{5\%}$, $N_{1\%}$). For the baseline execution, we set $N_{100} = 100$ (Pandora default setting) and the number of threads to 20. To compare the runtimes, we computed the speedups $S_{5\%} = \frac{T_{100}}{T_{5\%}}$ and $S_{1\%} = \frac{T_{100}}{T_{1\%}}$. A speedup below 1 indicates that the runtime with the convergence check enabled exceeds the runtime of computing all 100 replicates without checking for convergence. In cases where the bootstrap procedure does not converge before all 100 bootstraps are computed, the speedup is necessarily below 1 due to the additional convergence checks. To analyze the potential loss of accuracy of the early termination on the resulting stability analyses, we computed the deviation of PS values,

$$|PS_{5\%} - PS_{100}| \quad \text{and} \quad |PS_{1\%} - PS_{100}|,$$

as well as the deviation of PSVs,

$$|PSV^i_{5\%} - PSV^i_{100}| \quad \text{and} \quad |PSV^i_{1\%} - PSV^i_{100}|$$

for $i = 1, \ldots, M$.

Figures S8 and S9 show box plots of the speedup for PCA and MDS analyses for both convergence tolerance settings, respectively. Figure S10 shows box plots of PS and PSV deviations across all 13 datasets for PCA analyses. Figure S11 shows the PS and PSV deviations for MDS analyses. In all box plots, the dashed horizontal line in indicates the average value.

##### 2.4 Influence of $k$ on the PCS

Running Pandora on various population genetics datasets, we observed a substantial influence of the number of clusters $k$ used for $k$-means clustering on the PCS cluster similarity score. Figure S12 depicts this observation for the *HO-WE* dataset with 100 bootstrap replicates for $k \in [3, 15]$. While we do observe a general trend for a lower PCS with a higher number of clusters $k$, we also observe fluctuations for $k \in [5, 8]$ without a clear trend.

We thus strongly recommend providing a reasonable $k$ to Pandora based on prior knowledge or preliminary experiments, for instance, using the number of distinct populations in the dataset. We further note that the PCS should only be reported in conjunction with the number of clusters $k$.

#### 3 Software Versions and Hardware Setup

We used Pandora version 2.0.1 for all analyses. For PCA analyses within Pandora, we relied on *smartpca* version 8.

All analyses and benchmarks were run on two machines with a Xeon Platinum 8260 CPU (48 physical cores, 2.4 GHz, 754 GB RAM). For scripts and detailed instructions on how to reproduce all analyses, see `https://github.com/tschuelia/PandoraPaper`.

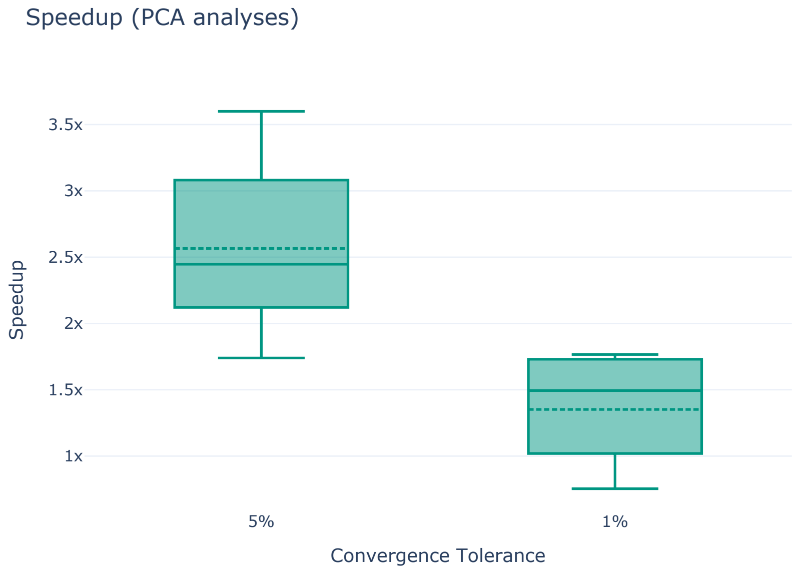


Figure S8: Speedup of the Pandora execution under the 5% and 1% convergence tolerance settings for PCA analyses. Note that we used 20 threads for all Pandora analyses.

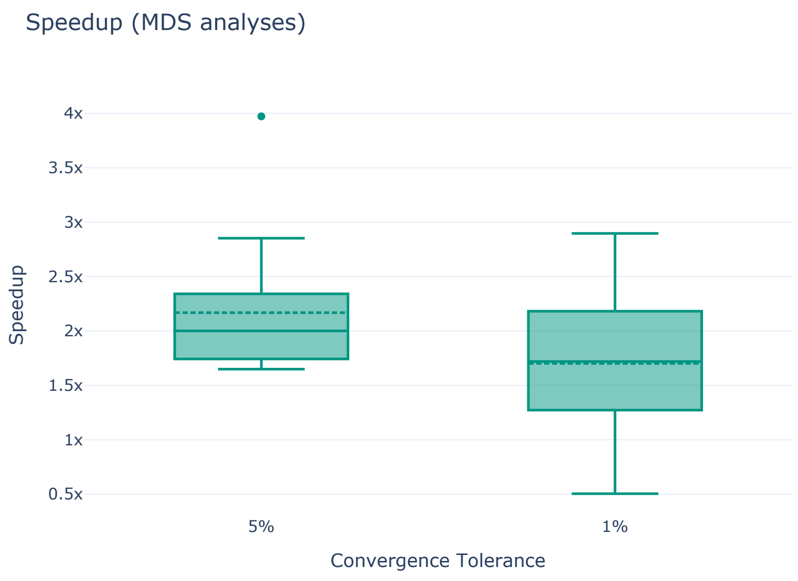


Figure S9: Speedup of the Pandora execution under the 5% and 1% convergence tolerance settings for MDS analyses. Note that we used 20 threads for all Pandora analyses.

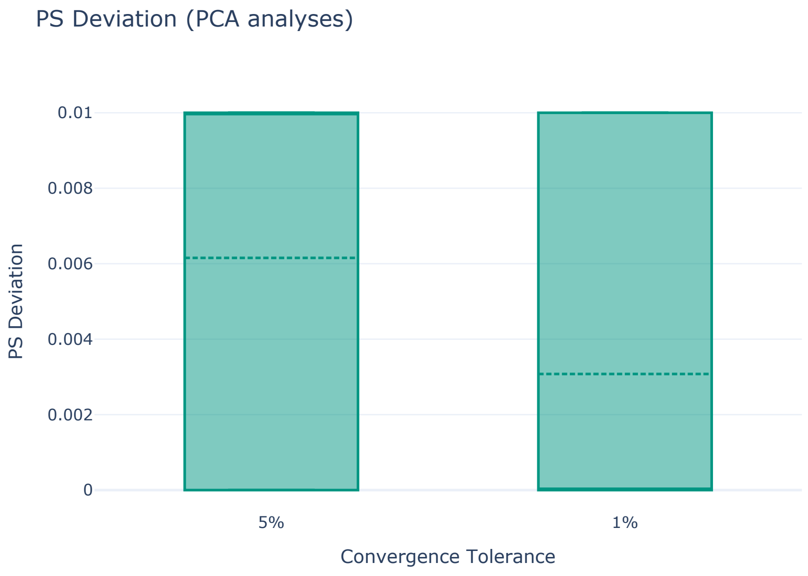


(a) Deviation of PS values.

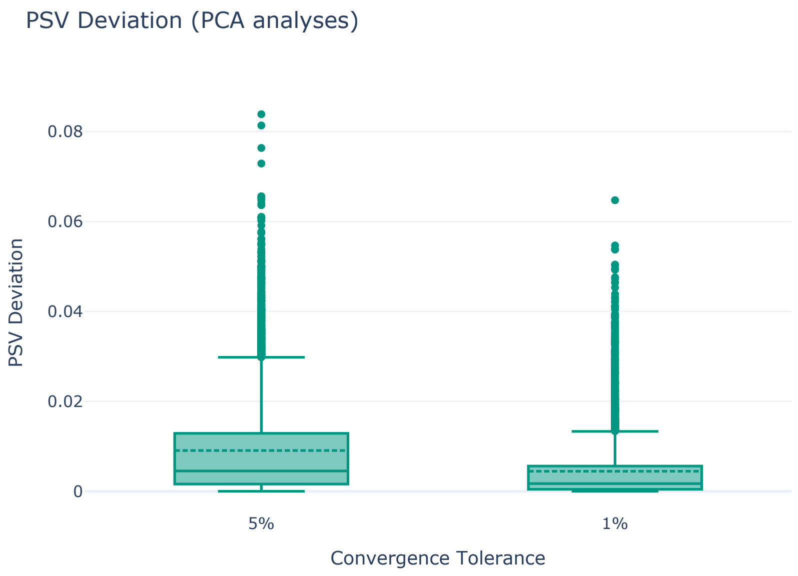


(b) PSV deviations for all individuals of all 13 simulated datasets.

Figure S10: Deviation of PS values and PSVs across all 13 simulated datasets under 5% and 1% convergence tolerance settings for PCA analyses

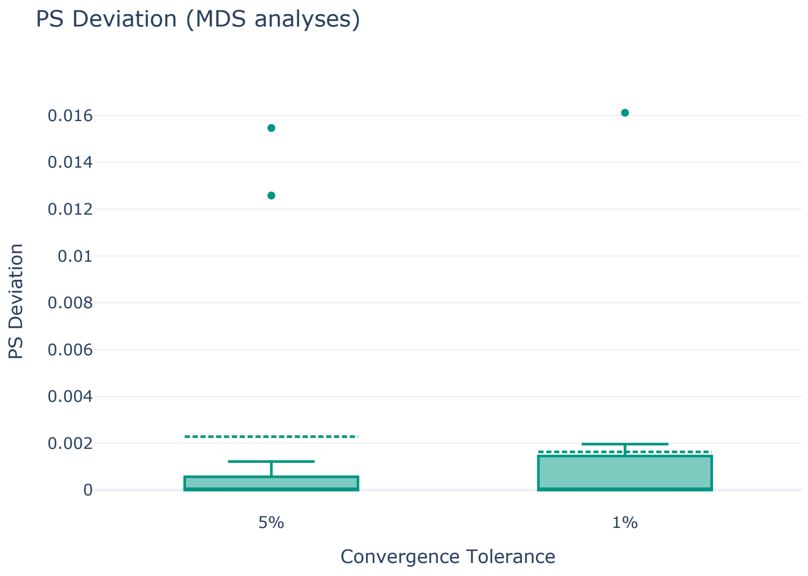


(a) Deviation of PS values.

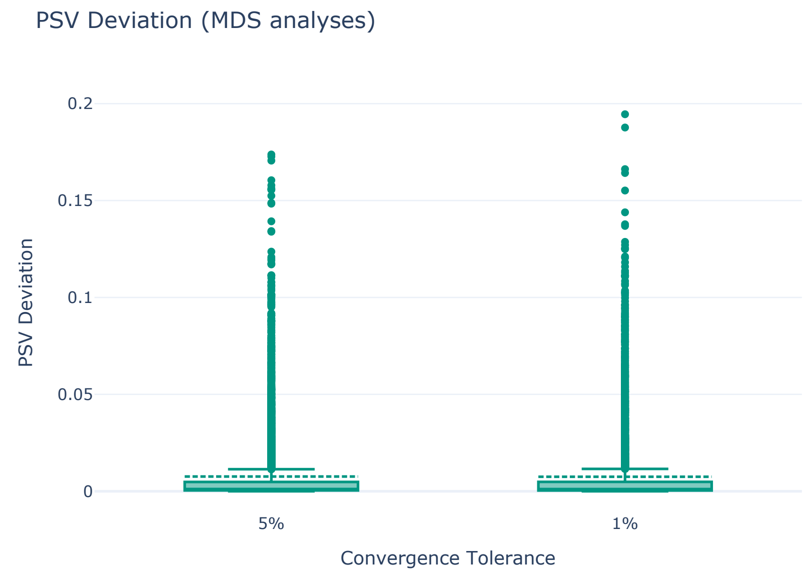


(b) PSV deviations for all individuals of all 13 simulated datasets.

Figure S11: Deviation of PS values and PSVs across all 13 simulated datasets under 5% and 1% convergence tolerance settings for MDS analyses

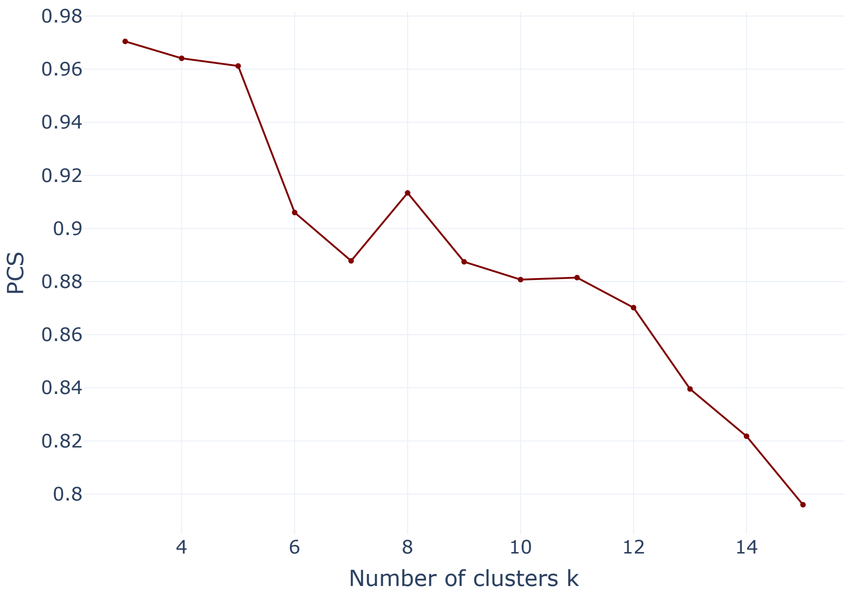


Figure S12: PCS for the *HO-WE* dataset as a function of the number of clusters $k$ used for $k$-means clustering.
